# Influence of gonadal and chromosomal sex on the brain transcriptome in a mouse species with natural sex reversal

**DOI:** 10.1101/2024.07.26.605342

**Authors:** Louise D. Heitzmann, Paul A. Saunders, Julie Perez, Pierre Boursot, Frederic Veyrunes

## Abstract

Sex chromosomes are expected to play a role in shaping the transcriptional architecture of sexual dimorphism, through the direct expression of sex-linked genes, by regulating autosomal genes, or in interactions with hormones. Yet, their degree of involvement remains elusive partly because chromosomal sex (XX/XY or ZZ/ZW) and gonadal sex (ovaries or testes) are usually inextricably intertwined. They are however dissociated in the African pygmy mouse, *Mus minutoides,* in which a feminizing X (X*) has evolved resulting in three female genotypes (XX, XX* and X*Y) and one male genotype (XY). Despite hormonal levels similar to the other females, X*Y females show distinctive phenotypes with greater fertility, divergent maternal care strategies and the masculinization of some traits (e.g. enhanced aggressiveness). By comparing the brain transcriptome of the four sexual genotypes, we show, here, that differential gene expression is mainly linked to gonadal sex (male vs. female) but also and significantly, to chromosomal sex, with expression patterns matching the singularity of X*Y female traits. Genes with such patterns are over-represented on sex and sex-linked chromosomes, and some are strong candidates to explain X*Y-specific behavioral and reproductive traits. We also report the preferential inactivation of the X* chromosome in XX* females, which could explain their trait similarities with XX females. Overall, we show that sex chromosomes have profoundly impacted the brain transcriptome in ways that reflect their new transmission modes and new resulting conflicts. This opens exciting prospects on the evolution of sex differences in relation to the dynamic of sex chromosome evolution.

## Introduction

Males and females differ for many traits (e.g., morphology, life history, behavior) as a result of the differential expression of underlying genes between the sexes (e.g. Rinn & Snyder, 2005; Pointer et al., 2013). Steroid hormones secreted by gonads and/or adrenal glands play a crucial role in mediating differences in gene expression and resulting phenotypes (Arnold, 2009; Van Nas et al., 2009; Phoenix, 2009; Trainor & Marler, 2002; Blencowe et al., 2022; Peterson et al., 2013). These influences are known as the organizational (i.e., perinatal) and activational (i.e., at adulthood) effects of hormones on sexual differentiation (Breedlove et al., 1999; Han & De Vries, 2003; Morris et al., 2004; Arnold, 2009; Konkle & McCarthy, 2011). In species with genetic sex determination, sex chromosomes (XX/XY or ZZ/ZW) are presumed to have a strong influence on sex differences as well (Rice, 1984; Mank, 2009; Dean & Mank, 2014). They are expected to be enriched in sex-biased genes (with different expression optima between the sexes) which can either directly influence sexually dimorphic phenotypes, or regulate the expression of other genes (autosomal or sex-linked) involved in differences between the sexes (Rice, 1984; Wijchers et al., 2010; San Roman et al., 2024).

Characterizing the impact of sex chromosomes on sexual differentiation is challenging because gonadal sex (hormones) and genotypic sex (sex chromosomes) are inextricably intertwined in most species with sex chromosomes. Nonetheless, they are decoupled in a few study systems, in which the relative influence of sex chromosomes and hormones on the transcriptional architecture of non-gonadal tissues was assessed (e.g. Blencowe et al., 2022; Ma et al., 2018). The study of genetically modified mice with sex-reversed XY females and XX males (Blencowe et al., 2022), or common frogs with naturally occurring sex-reversal (XX males; Ma et al., 2018) reported an underwhelming influence of the sex chromosome complement (XX vs. XY) on the establishment of transcriptomic differences between males and females. But given the particularities of these study systems -i.e., fixed genetic background of transgenic mice, and minimal sex chromosome differentiation as well as phenotypic variation between XX and XY males in common frogs-, the generalizability of the results is limited. A good model system to address this question requires two features: naturally occurring sex reversal, and phenotypic differences between individuals with the same gonadal but distinct chromosomal sex, such as found in a few rodent species (Saunders & Veyrunes 2021).

The African pygmy mouse, *Mus minutoides*, is an ideal system to study the role of sex chromosomes in the establishment of sex differences. This close relative of the house mouse naturally possesses three sex chromosomes: the classic X and Y, plus a feminizing X* chromosome (Veyrunes et al. 2010a). The latter originates from a still unknown feminizing mutation (∼1 My old) on the X chromosome that overrides the effect of *Sry* (Veyrunes et al., 2013). Consequently, there are three female genotypes: XX, XX*, X*Y, and one male genotype: XY. The X* chromosome is cytologically differentiated from the ancestral X with a different G-banding pattern and different size, and it stopped recombining with the X chromosome over a large chromosomal region (Veyrunes et al. 2010a, Baudat et al., 2019; unpublished data). Most interestingly, the emergence of the X* has led to drastic changes in the evolutionary trajectories of sex chromosomes (and potential sexual conflicts) in this mammalian species : the X* chromosome is female-specific and thus, supposedly free to reach a female phenotypic optimum; the Y chromosome which used to be male-specific is now shared with females; and the X chromosome now spends more time in males because 75% of females are X*Y (Saunders et al., 2022; Veyrunes et al., 2010a). Therefore, all of them are likely to influence sex-linked traits in a new way. Although adult X*Y females have functional gonads with a typical ovarian structure suggesting complete sex reversal (Rahmoun et al., 2014), they show divergent reproductive output and social/parental behaviors from both XX and XX* females, whereas the latter are remarkably comparable. X*Y females breed earlier and have a greater ovulation rate, which correlates with a larger average litter size (Saunders et al., 2014). They are more likely to be involved in uniparental care (while XX and XX* females are more likely to provide biparental care) and they have a better pup retrieval efficiency, but poorer nesting skills than the other females (Heitzmann et al., 2023). Finally, it has been shown that X*Y females display some traits considered to be typically male: enhanced territorial aggression, greater bite strength and low levels of anxiety assessed by behavioral investigations and measures of basal corticosterone levels (Saunders et al., 2016; Ginot et al., 2017; Veyrunes et al., 2024). All things considered, the behavioral and reproductive divergences observed among females imply differences in brain functioning (including the hypothalamo-pituitary axis regulating the reproductive cycle) and suggest differences in the sexual differentiation of the brain transcriptome. Because the three female genotypes have similar levels of sex steroid hormones (estradiol, testosterone), sex hormones and gonads are unlikely to be responsible to explain these differences, at least in adulthood (Veyrunes et al., 2024), suggesting a direct influence of sex chromosomes on brain gene expression and related phenotypes in *M. minutoides*.

In this study, we focus on the brain transcriptome of the four genotypes in the African pygmy mouse in order to understand the consequences of the emergence of the X* and sex reversal on gene expression profiles in the brain (i.e., brain transcriptome architecture). We aim to link these profiles with the different sexual phenotypes, and to assess the direct involvement of sex chromosomes in the development of sexual polymorphism (i.e., more than two sexual phenotypes in contrast to sexual dimorphism). We first compared overall gene expression in the whole brain of adult pygmy mice to evaluate the relative contributions of gonadal sex (i.e., male or female) and chromosomal sex (XX, XX*, X*Y and XY) to differences in gene expression. We then looked at the distribution of differentially expressed (DE) genes to assess the influence of sex chromosomes on gene expression (i.e., whether DE genes are over-represented on sex chromosomes; e.g., Khodursky et al., 2020; Ma et al., 2018; Ellegren & Parsch, 2007; Oliver & Parisi, 2004). Finally, we combined overall gene expression with allele-specific expression of genes showing allelic variation between the X and X* chromosomes to pinpoint potential sex-linked candidate genes at the basis of female phenotypic divergences.

## Results

### Differential gene expression

We sequenced the whole brain transcriptome of five adult *M. minutoides* of each genotypic sex (XX, XX*, X*Y females and XY males) bred in captivity. We detected 14,425 expressed genes, among which 1,362 are differentially expressed (DE) in at least one of the pairwise contrasts between the four sexual genotypes (log_2_ fold-changes for each contrast are presented in supplementary material S1). DE genes were clustered using a weighted correlation network analysis (WGCNA) based on their joint expression patterns across the four genotypes: 1,343 genes were split across eight clusters (thereafter ‘modules’) (fig. 1a), and the remaining 19 genes were not attributed to any module due to a low correlation with the module eigengene (i.e., a proxy of gene expression profiles inside a module). Most of the DE genes (∼78%) were grouped into modules that show a pattern of expression reflecting an influence of gonadal sex, i.e., males *vs* females. Among these, 756 genes (∼56% of DE genes) showed a male-biased expression pattern and were grouped into one module (MB module; fig1a, fig S1) while 308 genes (∼23% of DE genes) were mainly characterized by a female-biased expression pattern (FB module, fig1a, fig S1). They were however split into three modules due to different interactions with the sexual genotype (fig. 1a, fig. S1): female-biased but with variance between the three female genotypes (FB1 module, 19 genes), female-biased with greater expression levels in X*-carrying individuals (FB2 module, 276 genes), and female-biased with under expression in X*Y females (FB3 module, 13 genes). Patterns in the four remaining modules mainly reflected an influence of chromosomal rather than gonadal sex (20.5% of DE genes). Two of these modules reveal a significant impact of the presence of the Y chromosome (and/or heterogamety), with 67 genes overexpressed in individuals carrying a Y chromosome, i.e., both X*Y and XY individuals (Y-biased genes, YB module); and 145 genes underexpressed in Y-carriers (-YB module). A third module of 45 genes (X-dose module) reflects the number of X chromosomes an individual carries: gene expression was highest in XX females (two Xs), followed by XX* and XY individuals (one X), and then X*Y females (no X). Lastly, a module of 22 genes underexpressed in XX* and X*Y females indicates a negative correlation between the X* chromosome and gene expression (-X*B module).

**Figure 1.**
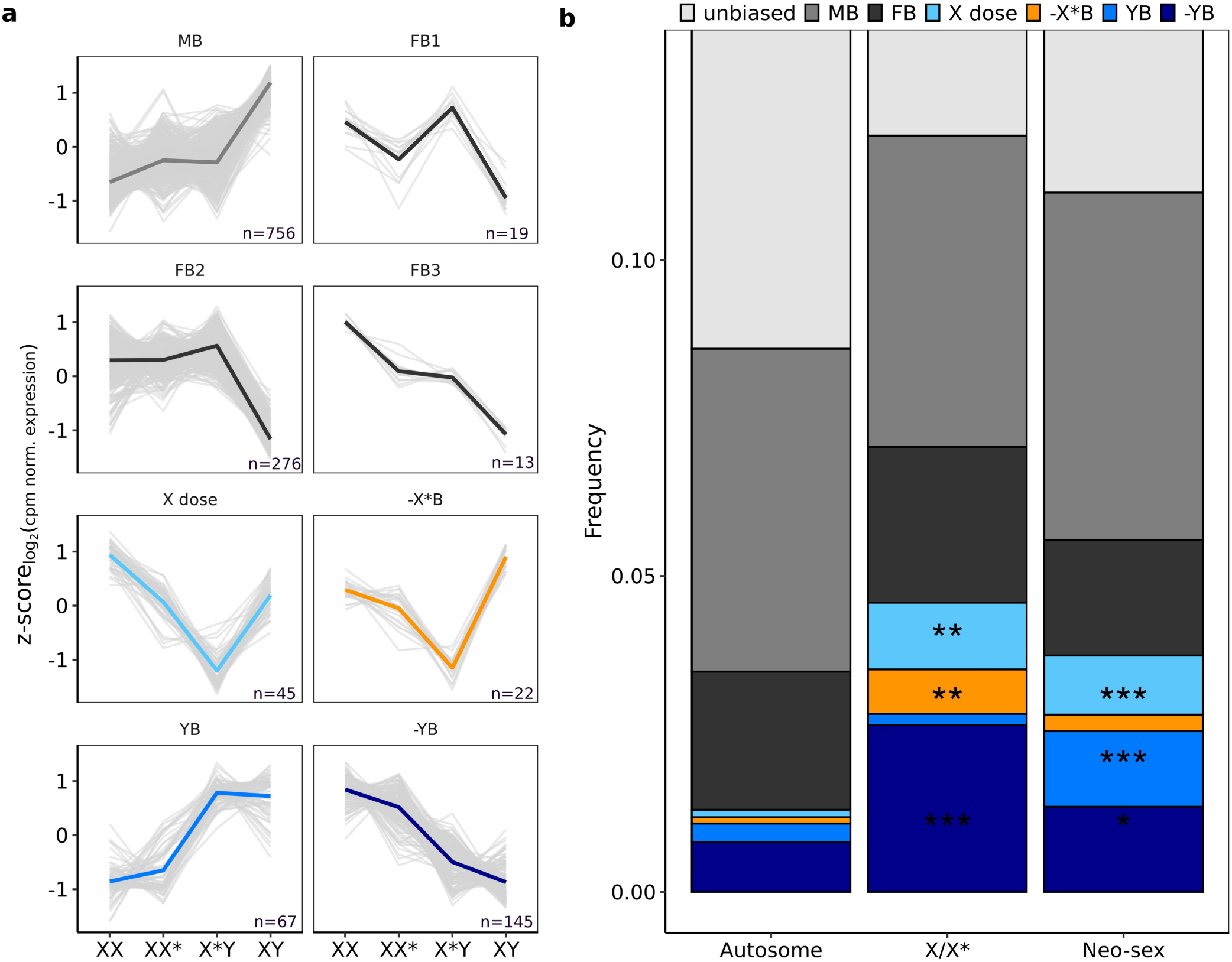
Brain transcriptional architecture in the African pygmy mouse. **a,** Clustering of the 1,362 differentially expressed genes in the brain of *M. minutoides*. Each graph corresponds to a module of genes with similar expression patterns identified by the WGCNA software. Light grey lines correspond to the average expression of each gene in the four genotypes. The average expression of all genes (i.e., representative of expression profile in a module) are represented by thick coloured lines for genes influenced by chromosomal sex and by thick lines in various shades of grey for genes influences by gonadal sex. Expression values are z-scores : cpm-normalized expression data are log_2_ transformed followed by scaling and centering. 19 genes were not assigned to any modules and are thus not represented. **b**, Distribution of unbiased and the 1,362 DE genes across the genome. Because sex chromosomes are known to be fused with autosomes in the African pygmy mouse, we separated the genome into three genomic compartment: autosomes, X/X* chromosomes and neo-sex chromosomes. In total, n_autosome_=11,015, _X/X*_=568,n_neo-sex_=2,674. Genes unassigned to a chromosome were not included in the diagram. Asterisks indicate the significance level of p-values from permutation tests (table S1), ***p<=0.001, **p<0.01, *p<0.05.

We next examined the distribution of DE genes of each module among genomic regions with different sex-linked transmission properties (autosomes and X/X* chromosomes). Because sex chromosomes are fused to autosomes in *M. minutoides* (producing neo-sex chromosomes with different patterns of inheritance; see details in methods; Veyrunes et al., 2004; Veyrunes et al., 2007; Baudat et al., 2019), we also included sex-linked autosomes into a single ‘neo-sex’ genomic region. We did not find an enrichment for genes with a difference in expression between males and females (FB and MB modules) in any of the three genomic compartments. However, genes in modules showing a genotype effect are non-randomly distributed, some are enriched on X/X* (permutation tests, X dose, p=0.007; -X*B, p=0.004; -YB, p=0.01) and/or on neo-sex chromosomes (permutation tests, X dose, p=0.001; YB, p=0.001, -YB, p=0.02) (fig. 1b, table S1).

### Gene ontology and function of DE genes

Male-biased genes are enriched in mitochondrial function GO terms, and female-biased genes are mainly enriched in neuron/axon development and synapse organization (see supplementary material S1). There is no significant enrichment of GO terms in modules showing a genotype effect. However, looking at ontologies gene-by-gene, we identified a few genes DE between females known to be involved in social behavior : *MaoA*, *Th*, *Sts* (Scott et al., 2008; Mahadevia et al., 2021; Scott et al., 2015; Mortaud et al., 2010), as well as in female reproductive cycle: *Amhr2 and Lhb* (Cimino et al., 2016; Huhtaniemi et al., 2006). We further found three genes DE between female genotypes that are known to escape X inactivation in *M. musculus*: *Kdm6a*, *Ddx3x* and *Eif2s3x* (Yang et al., 2010) (fig.2, table 1, supplementary material S1).

**Figure 2.**
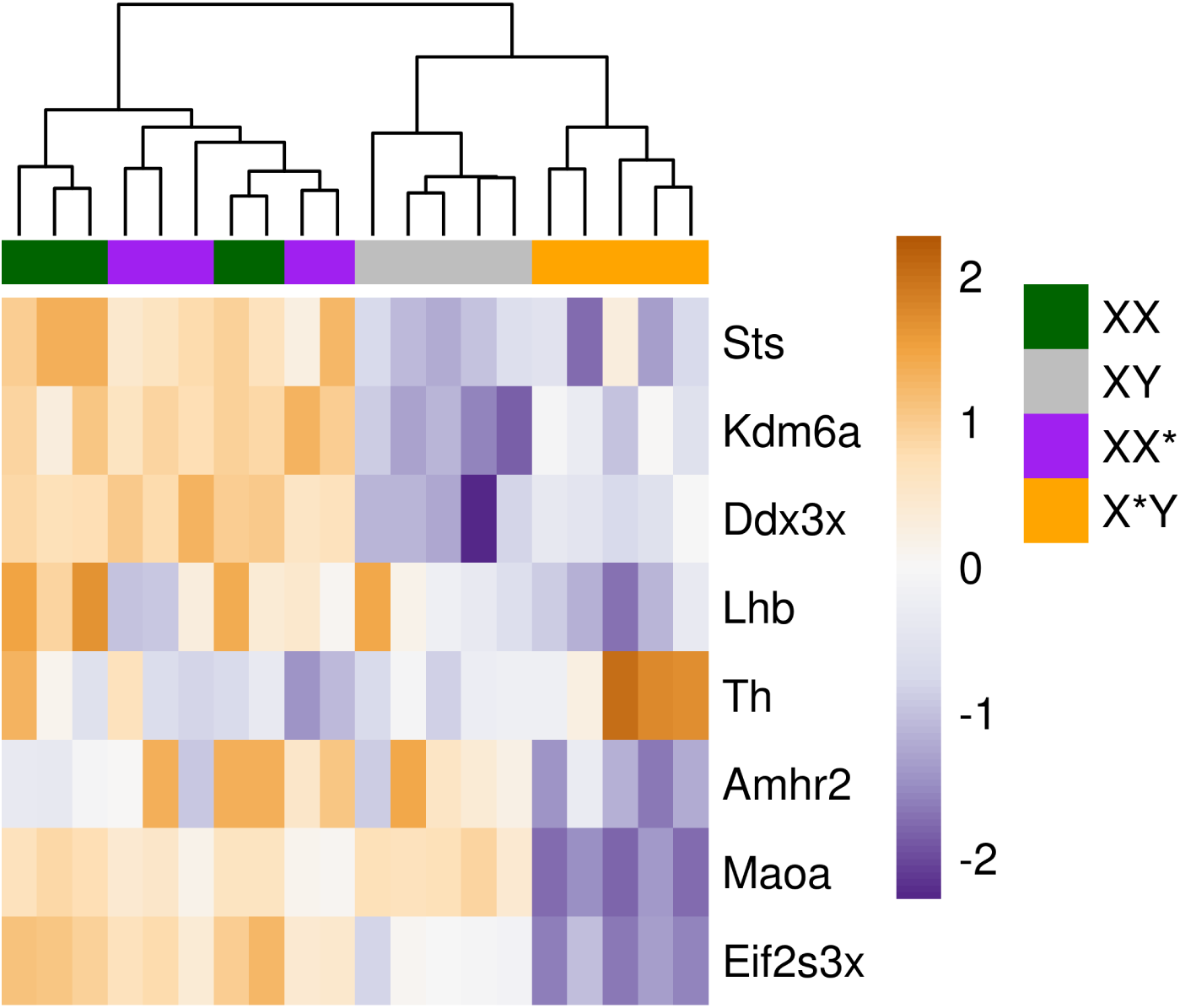
Heatmap of candidate genes expression. Values of relative expression are represented by z-scores.

**Table 1.**
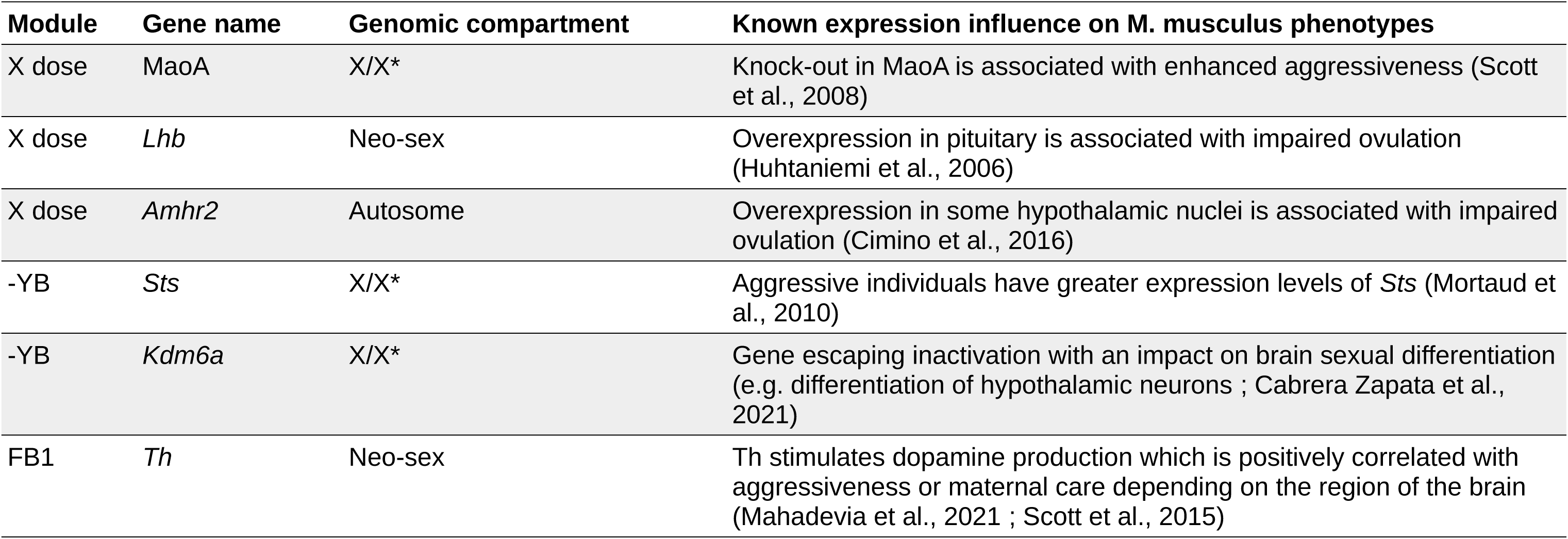
Candidate genes for phenotypic divergences highlighted in *M. minutoides,* based on *M. musculus* studies.

| Module | Gene name | Genomic compartment | Known expression influence on *M. musculus* phenotypes |
| --- | --- | --- | --- |
| X dose | *MaoA* | X/X* | Knock-out in *MaoA* is associated with enhanced aggressiveness (Scott et al., 2008) |
| X dose | *Lhb* | Neo-sex | Overexpression in pituitary is associated with impaired ovulation (Huhtaniemi et al., 2006) |
| X dose | *Amhr2* | Autosome | Overexpression in some hypothalamic nuclei is associated with impaired ovulation (Cimino et al., 2016) |
| -YB | *Sts* | X/X* | Aggressive individuals have greater expression levels of *Sts* (Mortaud et al., 2010) |
| -YB | *Kdm6a* | X/X* | Gene escaping inactivation with an impact on brain sexual differentiation (e.g. differentiation of hypothalamic neurons ; Cabrera Zapata et al., 2021) |
| FB1 | *Th* | Neo-sex | Th stimulates dopamine production which is positively correlated with aggressiveness or maternal care depending on the region of the brain (Mahadevia et al., 2021 ; Scott et al., 2015) |

### Differential expression between X and X* alleles : focus on XX* females

We were then interested in whether X/X*-linked genes showed a differential expression between X and X* copies, that would coincide with both the phenotypic distinctiveness of X*Y vs XX/XX* females and the similarities between XX and XX* females. We assessed allele-specific expression between the X and X* (see supplementary material for details on methods) using transcriptomic data of XX* females. Allele-specific expression could be tested for the 87 genes with fixed variants between the X and X*. We observe an overall bias towards higher expression of X alleles in all 5 females (two- to three-fold higher than X* alleles, fig. 3), with 77 genes (out of 87) showing a significant bias in that direction, and the remaining 10 showing no significant bias (supplementary material S2). This pattern is consistent across all individuals and regardless of the overall expression pattern of these genes, i.e., whether earlier detected as DE or not (fig. 3, supplementary file S2). Among these 77 genes, 68 were named in the genome annotation, which allowed us to retrieve their ontologies. There is no enrichment in GO terms for genes with a true allelic imbalance, but a gene-by-gene scan showed that many (∼50%) are involved in the development of the nervous system and RNA processing (e.g. transcription, splicing, methylation; see supplementary material S2). To identify whether this pattern of preferential expression is brain specific, we performed the same allele specific expression analysis using the kidney transcriptome. With the exception of one extreme individual that expressed only X* alleles, allelic expression was overall balanced between X and X* alleles (fig. S2-S3), as opposed to what is observed in the brain.

**Figure 3.**
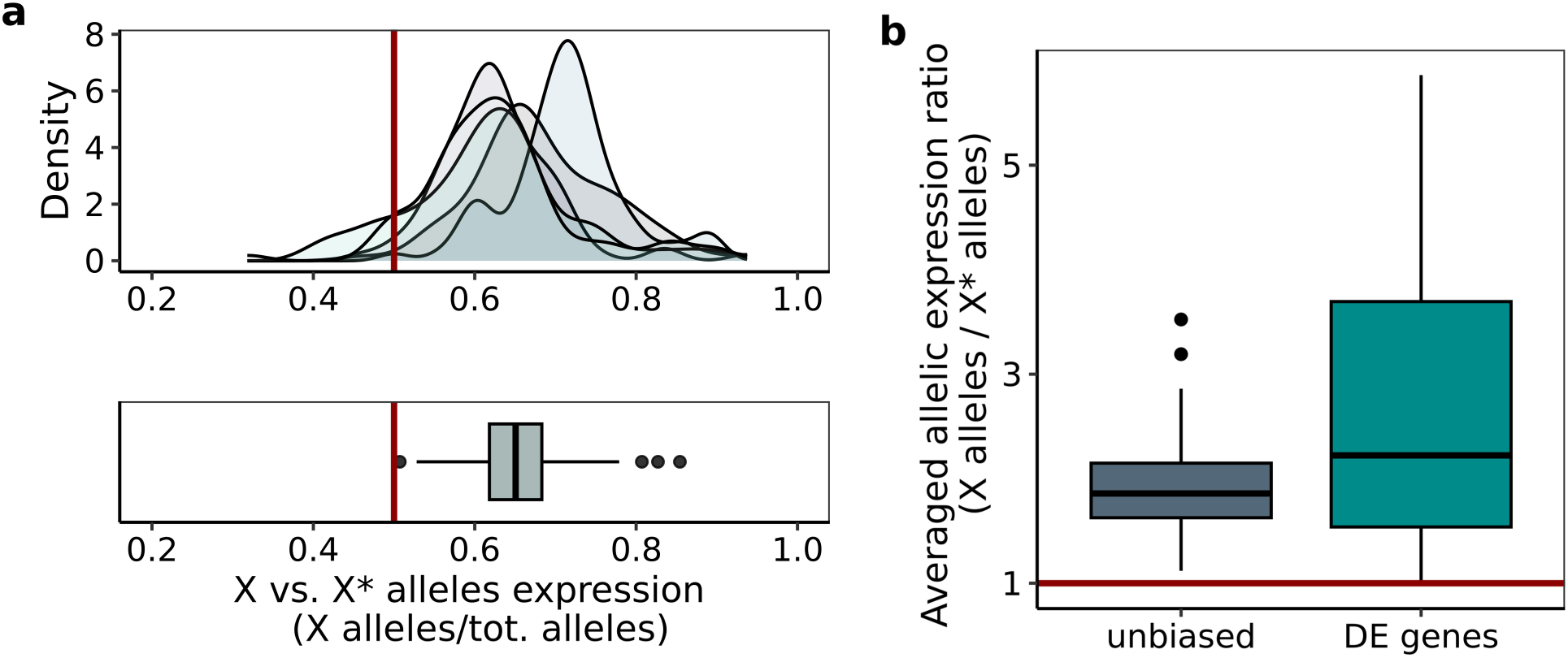
X/X*-linked allele-specific expression in the brain of XX*. **a,** Density distribution of X alleles in the five XX* females and average allelic expression ratio across the five females (boxplot below). Each shade ofcolour in the density plot correspond to a specific female. Values are a ratio of X alleles over all alleles per gene. The red line indicates a ratio of 0.5 and thus equal expression of both alleles. Values above 0.5 indicate a greater expression of X alleles and conversely, values below 0.5 indicate a greater expression of X* alleles. n_X/X*_=87. **b,** Averaged allelic expression ratios for unbiased and DE genes. Values in boxplots represents ratios of X alleles over X* alleles averaged across the five individuals, for each gene showing allelic variation, and according to their overall gene expression level: unbiased or DE. We removed one gene that was not detected in the overall gene expression analysis (i.e., did not pass the filter on minimal raw count reads kept for downstream analyses). The red line indicates a ratio of 1 with equal expression of both alleles. All values are above 1. On average, X alleles are expressed about twice as much (unbiased genes) or three times as much (DE genes) as X* alleles. Genes with true allelic imbalance are shown in supplementary material S2. n_unbiased_=76, n_DE genes_=10.

## Discussion

The African pygmy mouse has a complex sex determination system involving a dissociation between gonadal and chromosomal sexes: there are three female genotypes, including females carrying the Y chromosome. Previous studies in the species have shown that sex chromosome complement (XX/XX*/X*Y) influences various life history and behavioral traits in females despite complete sex reversal and similar hormonal levels, thus revealing the role of sex chromosomes in shaping sex-linked phenotypes. Although a majority of genes displaying differential expression show gonadal sex-linked expression, our results highlight a significant influence of sex chromosome complement on gene expression as well. Moreover, genes influenced by chromosomal sex are over-represented on sex and neo-sex chromosomes (fig. 2b): these regions have a decisive influence on the brain transcriptional architecture and are thereby an important driver of phenotypic divergence between female genotypes. Finally, allele-specific expression analyses revealed a preferential inactivation of the maternal X* chromosome in the brain of XX* females. This striking result is concordant with the recurring observations of comparable phenotypes in XX* and XX females (Saunders et al., 2014; Saunders et al., 2016; Heitzmann et al., 2023; Veyrunes et al., 2024).

### a) The brain transcriptome reflects an influence of the sexual genotypes

Brain transcriptome analysis reveals that 9.4% of all expressed genes (1,362 out of 14,425) are differentially expressed (DE) between at least two of the four sexual genotypes (XX, XX*, X*Y and XY). Most of these genes (1,064 genes, i.e., ∼78%) group females together against males, which is consistent with a transcriptional architecture that mainly reflects gonadal sex, and therefore a strong influence of sex hormones. Nonetheless, a remarkably high proportion of DE genes (∼20.5%) is predominantly influenced by the sex chromosome complement, with X*Y females grouped together with males or opposed to all genotypes. In comparison, in common frogs (*Rana temporaria)*, which also present naturally occurring sex-reversed individuals (XX males), a whole-body transcriptome study showed that only 0.06 % of DE genes showed a genotype effect, with gonadal sex being the main factor influencing variance in gene expression (Ma et al., 2018). Recently, the same result was found using liver and adipose tissue transcriptomes in a genetically modified mouse model with sex-reversed XY females (lacking *Sry)* and XX males (with an *Sry* transgene) (Blencowe et al., 2022), where only ∼0.01-0.02 % of DE genes were found to be influenced by chromosomal sex (∼10 genes whose expression is influenced by the genotype in comparison to ∼500-1000 DE genes influenced by gonadal sex). These contrasting results most likely reflect the discrepancies between the sporadic, instantaneous cases of sex-reversal (environmentally and/or genetically induced) and the novel sex determination system in the African pygmy mouse that emerged around 1 Mya, involving a new genomic architecture : three differentiated sex chromosomes (and associated neo-sex chromosomes), with different ancestries and inheritance patterns (Veyrunes et al. 2013), leaving time for gene expression to evolve accordingly. All things considered, our results illustrate that within a same hormonal background and gonadal sex (male or female), sex chromosomes and neo-sex chromosomes participate in shaping the brain transcriptome and sexual polymorphism in *M. minutoides.* It would thus be interesting to expand this research and include other systems with atypical sex determination system (e.g. species lacking the Y chromosome, species with both female/male heterogamety; see Saunders & Veyrunes, 2021) to further our understanding of how sex chromosomes can influence gene expression and resulting phenotypes.

Finally, it is interesting to note that the brain transcriptome of X*Y females shows signs of masculinization (expression levels similar to those of males), which is consistent with some of their behavioral traits considered to be male-typical (e.g. enhanced territorial aggression; Saunders et al., 2016). Similarly in the wild turkey (*Meleagris gallopavo*), males exhibit dominant or subordinate phenotypes, which involves variance in the degree of sexual dimorphism and correlates with the magnitude of sex-biased gene expression (Pointer et al., 2013). Therefore, while it is often complicated to test the relationship between sex-biased gene expression and sexual dimorphism because the latter is often envisaged as dichotomous, we highlight that species such as in *M. minutoides* or the wild turkey (Pointer et al., 2013) displaying sexual polymorphism are highly valuable.

### b) Sex chromosome contribution to variation in gene expression

Theoretical models predict that sex chromosomes should play a large role in shaping the transcriptional architecture of sexual dimorphism due to an enrichment in sex-biased genes (e.g. Rice, 1984). Although a few empirical studies support these models (e.g. Zemp et al., 2017; Kottler & Schartl, 2018), the sex-linked genetic bases of sexual dimorphism are still poorly documented and understood. First, there is a serious lack of data that link sex-biased expression to sexually dimorphic phenotypes (Mank, 2017, Dean & Mank, 2014). Then, evidence for the direct involvement of sex chromosomes is hampered by the fact that their effects are often confounded with those of sex hormones (i.e., chromosomal sex matches gonadal sex). In fact, their effect has even been neglected to the benefit of sex hormones when considering sex differences in brain and behaviors. Therefore, sex chromosomes’ influence on gene expression, and their implication on sexual dimorphism remains debated or even overlooked depending on studies. Here, we have a system with two hormonal types but more than two sexual genotypes and highlight a direct influence of the sex chromosome complement on gene expression. Approximately three quarters (212 genes) of the DE genes whose expression varies with genotypic sex display identical ex-pression levels in X*Y and XY individuals *vs.* XX and XX* females (YB and -YB genes, Fig.1a). They are found across the whole genome, but are over-represented on X/X* and neo-sex chromosomes (Fig.1b), suggesting that their expression is under the influence of the Y (and neo-Y) and/or the number of X/X* chromosomes (including epistatic interactions between these chromosomes). The remaining quarter of DE genes with an expression influenced by genotypic sex (67 genes), have different expression in X*Y females *vs.* all other genotypes, suggesting an X* and/or an X chromosome effect (e.g. X dose), with different impacts on gene expression between the X and X* chromosomes (fig 1b). These genes are mainly over-represented on X/X* chromosomes and because the X* chromosome stopped recombining with the X chromosome over a large region of the chromosome, it is very likely that some, if not most of them, are in the non-recombining region and therefore, could have diverged following the emergence of the X*. Future investigations including an outgroup to orientate changes in gene expression could help identify the evolutionary mechanism driving sex-biased gene expression in *M.minutoides.* All things considered, X*Y females display a singular transcriptomic architecture, that either distinguishes them from the other females, or from all the other genotypes, in line with phenotypic observations. These results illustrate that the emergence of the X* chromosome has profoundly altered the evolution of sex-linked regions at the expression level. The new inheritance patterns of sex chromosomes have changed their evolutionary trajectories: the female-specific X*, the biparentally inherited Y, and the X that now spends more time in a male than in a female context (Saunders et al., 2022) drive the evolution of differential expression between sexual genotypes and most likely, sexual polymorphism in brain and behaviors. Consequently, sex-linked regions (the X, X* and Y chromosomes and neo-sex chromosomes) act in orchestra to shape the brain transcriptome of *M. minutoides* and resulting sexual phenotypes. This makes the African pygmy mouse an excellent study model to investigate the dynamics of gene expression and the evolution of sexual dimorphism in relation to sex chromosomes’ own dynamics and inheritance pattern.

### c) Sex-linked candidate genes for the molecular bases of the different female phenotypes

Previous studies showed that X*Y females differ from both XX and XX* females along many reproductive and behavioral traits (Saunders et al., 2014; Saunders et al., 2016; Heitzmann et al., 2023). Some of the sex-linked genes found to be DE in this study could be responsible for these differences and are thus worth investigating further. For instance, the Luteinizing hormone subunit beta (*Lhb;* neo-sex-linked) is essential for gonadal functions and its overexpression implies an increased activity of LH, which is associated with impaired ovulation and follicle maturation (Coss, 2018; Huhtaniemi et al., 2006). *Lhb* is underexpressed in X*Y females in comparison to XX and XX* females, which is consistent with the fact that X*Y females breed earlier and have an ovulation rate 1.5x greater than the other females (Saunders et al. 2014). Interestingly, another gene substantial for reproductive functions, the Anti-Mullerian Hormone Receptor 2 (*Amhr2*; autosomal), whose overexpression alters ovulation (Cimino et al., 2016), follows the same expression pattern as *Lhb*, suggesting that it is influenced by X-linked genes dosage (with different effects between the X and X*) and could be involved as well.

We also identified two sex-linked genes known to regulate aggressiveness: monoamine oxydase A (*MaoA;* X-linked), and Steroid sulfatase (*Sts,* sex-linked) (Mortaud et al., 2010; Scott et al., 2008), with potential roles in the greater aggressiveness levels observed in X*Y females (Saunders et al. 2016). Both are found to be underexpressed in X*Y females when compared to the other females. Sts does not, however, fit the observed phenotypes since its over-rather than underexpression has been linked to aggressive behaviors (Mortaud et al., 2010). In contrast, *MaoA*, an enzyme involved in dopamine degradation, is a strong candidate to explain phenotypic differences, as its down-regulation has been related to enhanced aggressiveness in mice (Scott et al., 2008). Moreover, we found another DE gene involved in the dopaminergic system, Tyrosine hydroxylase (*Th,* the rate limiting enzyme for dopamine synthesis; neo-sex-linked), known to stimulate both aggressive and maternal behaviors in laboratory mice (Mahadevia et al., 2021; Scott et al., 2015). *Th* is significantly overexpressed in X*Y females when compared to XX, XX* and XY individuals (see supplemetary material S1). Taken together, our results highlight two sex-linked genes from the dopaminergic pathway that are strong candidates to correlate with divergences in social and parental behavior in *M. minutoides*.

It is also worth noting that we found the lysine demethylase 6a gene *Kdm6a* to be underexpressed in X*Y females. *Kdm6a* is X-linked and escapes X chromosome inactivation (XCI) in laboratory mice (Yang et al., 2010). It has also been shown to up-regulate *Xist* expression driving X inactivation (Lin et al., 2023) and to regulate neuron sexual differentiation in the ventromedial hypothalamus (Cabrera Zapata et al., 2021) as well as the paraventricular nucleus of the hypothalamus (Xu et al., 2008). These regions of the brain are both involved in sex differences in social, reproductive and parental behaviors (Bayless & Shah, 2016; Scott et al., 2015; Hashikawa et al., 2017; Musatov et al., 2006). Here, XX and XX* females have a greater expression level of *Kdm6a* when compared to X*Y and XY individuals. This first suggests that *Kdm6a* also escapes inactivation in *M. minutoides,* and that it could regulate the expression of genes involved in differences in social/parental behaviors between female genotypes (Saunders et al., 2016; Heitzmann et al., 2023). On that note, we confirmed that *Ddx3x* and *Eif2s3x* seem to escape inactivation in *M. minutoides,* as they do in *M. musculus,* with greater expression levels in XX and XX* females. Nonetheless, the expression of a single copy of these genes does not seem to alter female development in contrast to observations on females with Turner syndrome (i.e., 45, X0 females; see Berletch et al., 2010). Further comprehensive studies with RT-qPCR and *in situ* hybridization in specific regions of the brain, such as the hypothalamus, may provide more insights into the genetic basis of the sexual phenotypes in the African pygmy mouse.

### d) Allele-specific expression and X/X* chromosome inactivation in XX* females

Based on the identification of fixed sequence differences between X and X* chromosomes, we were able to assess allele-specific expression of those chromosomes. Looking at the brain of XX* females that presumably expresses both alleles, we show that XX* females preferentially expressed X alleles (fig. 3). This pattern is seen for all genes (unbiased and DE) and consistent across individuals, which suggests a chromosome-wide effect and thus, the preferential inactivation of the X* chromosome (X*CI) in the brain of XX* females. This could explain similar overall gene expression patterns found here between XX* and XX females, and by extension their similar behavioral and fertility phenotypes (Saunders et al., 2014; 2016; Heitzmann et al., 2023). We note though that none of the genes with documented allelic imbalance is a known candidate to be involved in female phenotypes. Finally, because there is an overall balanced expression of X/X* alleles in the kidney transcriptome (fig. S2-S3), in line with previous observations of random inactivation of X and X* observed in embryo fibroblast cultures (Veyrunes & Perez, 2018), the preferential inactivation of the X* chromosome could be brain specific. This observation is intriguing, because it contrasts with the assumption of overall random XCI in eutherian mammals. In fact, a very limited number of studies have reported evidence for a skewed XCI, but it mainly involves the inactivation of the paternal X, as shown in Marsupials (Cooper et al., 1993) or in the brain of neonatal mice (Wang et al., 2010). Hence, to our knowledge, our results appear to be the first instance of the preferential inactivation of the maternal X in a mammalian species. In the future, it could be worth investigating genes involved in X chromosome inactivation (e.g. *Xist* or its antisense gene, *Tsix,* unfortunately not annotated in our genome) including polymorphism in their regulatory regions (e.g. Hudson, 2003; Jiang, 2010; Bongiorni et al., 2014), to understand the proximate mechanism behind this genomic imprint.

In the following, we discuss a few hypotheses that could explain X* preferential inactivation. First, the non-recombining part of the X* chromosome is female specific. In absence of recombination, the X* could accumulate mutations in cis-regulatory elements that have deleterious effects on the level of expression (Lenormand et al., 2020). However, in X*Y females, the X* is hemizygous and faces an already degenerated Y chromosome, exposing it to a strong purifying selection. Hence, the X* is unlikely to degenerate, or at most to a very limited extent. Consistent with this reasoning, we do not find evidence for the preferential expression of X alleles in the kidney (fig. S2-S3), which strongly suggests that skewed X*CI in the brain of XX* females is not a response to X* chromosome degeneration.

On the other hand, we previously mentioned that the emergence of the X* has led to new evolutionary trajectories of all sex-linked regions, which might have laid ground for new sexual conflicts and influence how genes involved cope with them. For example, because the X* is female-specific (i.e., transmitted exclusively from mothers to daughters), gene expression should be free to evolve towards a female phenotypical optimum. On the other hand, the X chromosome now spends more time in a male environment in comparison to what prevails in an XX/XY system (Saunders et al., 2022), which may induce X chromosome masculinization. These new evolutionary trajectories of X and X* could drive new genomic conflicts with divergent interests between chromosomes. It is also noteworthy that the X chromosome is enriched in genes involved in brain functions (e.g. neurodevelopment, behaviors; Skuse, 2006) and because the brain regulates many reproductive phenotypes, from behaviors to reproductive cycle, both are likely to be prime targets of sexual conflicts in relation to divergent phenotypic interests between the sexes.

It is however puzzling that the X* rather than the X is preferentially inactivated in XX* females. Under the predictions of X masculinization and X* feminization, we could expect preferential inactivation of the X in XX* females. It should be noted, however, that this is no longer true if the X* is beneficial for females only when it is accompanied by the Y chromosome, with which it co-evolves since its emergence. This would imply epistatic interactions between X* and Y chromosomes in X*Y females. Whether this interaction is cooperative (i.e., both contributing to make better females) or antagonistic (e.g. the Y still contributing to preserve male quality) is not known but in either case, X* without Y could be dysfunctional and selection would thus favor its silencing, at least in the brain. Under this assumption, we could predict the preferential inactivation of the X* in gonads as well, the most sexually dimorphic organs (e.g. where Y-linked genes expression in ovaries would be deleterious). Therefore, further investigations on the gonad transcriptome could help understand the ultimate cause behind the preferential expression of the paternal X in XX* females in the brain of the African pygmy mouse.

## Conclusion

The African pygmy mouse is a peculiar species with three sex chromosomes: the classical X and Y, and a feminizing X* chromosome. In this study, we have demonstrated that the new mode of transmission of sex chromosomes profoundly affects the brain transcriptome, which reflects females’ phenotypic divergences and the third sexual phenotype highlighted in previous studies (Saunders et al., 2014; Saunders et al., 2016; Heitzmann et al., 2023; Veyrunes et al., 2024). While it is often difficult to establish a link between sexual dimorphism, sex-biased expression and a direct role of sex chromosomes, we were able to find several sex-linked candidate genes that are worth investigating further: their expression pattern varies along with the genotype and could underlie the different female phenotypes, directly and indirectly related to the brain (e.g. maternal behaviors, ovulation).

Finally, this is the first study to our knowledge reporting preferential inactivation of the maternal X chromosome (X* in our case) in a mammalian species, which seems to be brain specific and could explain why XX* females are more similar to XX females. Our results emphasize that the evolution of this unique sex chromosome system in the African pygmy mouse may have opened new avenues to sexual phenotypes, reflecting new sexual conflicts. In the future, looking at other organs and including an outgroup species could provide insights into the evolutionary mechanism that drive the evolution of gene expression and sexual polymorphism in *M. minutoides*.

## Material & Methods

### a) Animals and sampling

Animals were bred in our breeding colony at the university of Montpellier (CECEMA facilities). The colony was established from wild caught animals in the Caledon Nature Reserve, in South Africa (Veyrunes et al., 2010a; see Heitzmann et al,. 2023 for further details on rearing conditions). In this study, we sampled the whole brain of 20 virgin individuals (5 per genotype: XX, XX*, X*Y and XY) aged between 5-9 months. Females were genotyped by PCR of *Sry* using gDNA extracted from tail tip biopsies following Veyrunes et al. (2010a) or by karyotyping using bone marrow of yeast-stimulated individuals (see Veyrunes et al., 2010b). We used unrelated individuals when possible, but some related animals (siblings) were included both within and between genotypic groups. Mice were euthanized by cervical dislocation and whole brains were collected and preserved in 2 mL tubes with RNAlater® (ThermoFisher). Samples were maintained at 4°C for 48 hours and then stored at −80°C until processing. Whole brain RNA was extracted under a sterile environment following RNeasy® Plus Mini Kit (Qiagen) protocol for animal tissue, with three steps: tissue disruption, homogenization and RNA purification. Brains were grounded into a fine powder in liquid nitrogen using a pestle and a mortar to assure cell membrane disruption. Tissues were then homogenized with a lysis buffer consisting of 24 μL of β-mercapto added to 2400 μL of RLT Plus (4x volume recommendations for 20-30 mg of tissue). After homogenization, we split the lysate into four tubes to prevent clogging of the columns during RNA purification. We purified two tubes per sample following the manufacturer’s provided protocol and stored the other two at −80°C. Total RNA was eluted in 50 μL of RNase-free water and stored at −80°C until further use.

### b) Sequencing and mapping

Stranded RNA-seq libraries preparation and cDNA sequencing were performed by Fasteris SA (Switzerland) using an Illumina HiSeq 3000/4000. Sequencing generated ∼13-20 million of 150 pb paired-end reads per sample. We assessed quality control of the samples using FastQC v0.11.9. Reads were filtered and trimmed using cutadapt v2.8 to keep reads with a minimal quality of 30, a minimal length of 75 pb and to remove adapters as well as N flanking bases. Trimmed reads were then mapped on XX female reference genome (Mus_minutoides_I2396, GenBank accession number GCA_902729485.2) using STAR software v2.7.10b. We used the XX genome masked at fixed variable sites between X and X* chromosomes to account for potential mapping biases among genotypes (see supplementary methods). We allowed for a maximum of 6 mismatches in 150 bp paired-end reads (option --outFilterMismatchNoverLmax 0.02), flagged duplicate and multi-mapped reads (option --bamRemoveDuplicates Type UniqueIdentical) and sorted reads by coordinates. We then performed gene quantification using the featureCounts software v2.0 from the subread package (Liao et al., 2013) for reverse stranded data, at the exon level and with a minimum mapping quality score of 10. The latter returned one text file per sample, including read counts for each gene annotated on the XX reference genome (Ensembl, Mus minutoides-GCA_902729485.2-2023_03). For some genes, symbol and names were not present in the ensembl annotation file, gene names were thus retrieved (when unambiguous, by considering single/consensus hits, coverage and identity percentage >80%) using the nucleotide blastn tool from ncbi (https://blast.ncbi.nlm.nih.gov/Blast.cgi).

### c) Differential expression analysis

Read counts were filtered to keep genes with a minimal coverage of 10 reads in at least 5 individuals (i.e., size of each genotypic group) and subsequent gene expression analysis was performed using EdgeR on R v4.3.2 (R core Team, 2023). Library sizes were normalized (TMM method) to account for differences in sequencing depth between samples and to compare gene expression across genotypes. We made multiple pairwise comparisons between the four genotypes (i.e., 6 contrasts, e.g. XX *vs* X*Y) to detect genes that were differentially expressed between at least one contrast, with a Benjamini-Hochberg correction for multiple testing. Genes with a fold-change value greater or equal 1.5, and with a false discovery rate (FDR) below or equal 5% were considered as differentially expressed (DE). Genes not detected as DE are referred as ‘unbiased’ throughout the paper. All genes (unbiased and DE) in *M. minutoides* were log normalized for sequencing depth using the cpm function in EdgeR, which incorporates the normalized library size. We further scaled and centered the expression of each gene to a mean of 0, and a standard deviation of 1 across samples (referred to as z-score values) for graphical representations of DE genes across genotypes.

### d) Gene expression clustering

DE genes were grouped according to their expression pattern by an unsupervised hierarchical clustering method using the Weighted Gene Correlation Network Analysis (WGCNA) software v.1.72 (Langfelder & Horvath, 2008). WGCNA clusters genes whose expression are highly correlated (i.e., co-expressed genes) into modules. We used a bidweight midcorrelation with a maximum percentile outlier of 0.05 following authors’ recommendations. We chose a soft thresholding power of 13 based on the scale-free topology fit indices (R² values > 0.9 and slope close to 1) to detect ‘signed’ modules of co-expressed genes (signed modules distinguish between mirror modules, with only positively correlated genes). We allowed for a minimal module size of 5 genes and merged modules with a threshold height of 0.24 on the clustering dendrogram (i.e., similar modules). Using the module-biological trait correlation function of WGCNA, each module was related to a biological category: gonadal sex (male versus female) and genotypic sex with three categories: presence of the Y chromosome, the X*, and number of X chromosomes (fig. S1).

### e) Gene assignment to chromosomes and DE genes distribution

Genes in the *M. minutoides* genome annotation file are assigned to a genomic scaffold and not a chromosome. Thus, to position each gene on a specific chromosome, we aligned each scaffold on *M. musculus* reference genome GRCm39 using LastZ v1.04.22 with the following parameters: --step=20 --ambiguous=iupac --seed=12of19 --hspthresh=3000 --gappedthresh=3000 --gfextend --chain --gapped --inner=2200.

Alignments were filtered to keep reciprocal best alignments only using the chain/net procedure (scripts downloaded from https://github.com/ENCODE-DCC/kentUtils/tree/master/src/hg/mouseStuff; Kent et al., 2003), and further filtered manually (min. alignment score : 300,000, min. percentage of query sequence involved in alignment: 5%). Scaffolds that did not anchor were defined as unassigned genomic regions. Then, we merged our annotation file with the alignment output and corrected for chromosome homologies between *M. musculus* and *M. minutoides* to assign each gene to a chromosome in our species (Veyrunes et al., 2006).

For following analyses, we distinguished three different gene categories, linked to different genomic compartments of interest: (i) genes carried by the X (and therefore X*) chromosome (568 genes), (ii) genes with autosomal transmission (11,0115), and genes linked to neo-sex chromosomes (*i.e.*, autosomes fused to sex chromosomes in *Mus minutoides*; which have therefore acquired a sex-linked transmission; Veyrunes et al., 2004; Veyrunes et al., 2007; Baudat et al., 2019; genes found on chromosomes 1, 13 and 16: 2,674 genes).

To test whether the sex chromosomes (X/X*) or neo-sex chromosomes are enriched in DE genes, we used Pearson χ^2^ (one-sided hypothesis) and a permutation test. For the latter, we sampled n genes from the genome (n=number of genes in that compartment) 1000 times, and compared the proportion of DE genes in the different modules in each permuted dataset to the observed proportions. P-values were determined by the number of times expected frequencies were greater or even to observed frequencies over the 1000 iterations. Genomic regions enriched in DE genes identified by both methods (χ^2^ and permutations) were considered truly enriched. P-values from permutation tests were reported in the main text as they are more stringent.

### f) Sex-specific allelic expression

We focused on allele-specific expression on X/X*-linked genes to identify differences of expression between the X and X* chromosome that could underlie phenotypic differences between females. X/X*-linked fixed variants were identified by SNP calling using transcriptomic data (see supplementary methods). Allelic read counts were retrieved using the ASEReadCounter tool from GATK v4.4.0 and filtered for mapping quality (≥ 10), base quality (≥ 10) and the minimum number of bases that passed filters (≥ 10). The ASEReadCounter tool returns allelic counts at the SNP level, we therefore retrieved allelic counts at the gene level using the ASEReadCounter* pipeline (Mendelevich et al., 2020), which interpolates SNPs and genes’ position on genomic scaffolds. We filtered genes with a minimal count of 10 reads in at least 5 allelic counts out of the 10 (2 alleles x 5 sample replicates). When an X* allele was found in XX females or conversely, when an X allele was found in X*Y females, we removed the gene involved (i.e., misdiagnosed SNP: shared polymorphism between chromosomes). We modelled allele-specific expression fitting a negative-binomial distribution, using gene ID and alleles’ origin as fixed effects and with an interaction term. Post-hoc pairwise comparisons between reference (X) and alternate (X*) alleles for each gene were made applying a Benjamini-Hochberg correction for multiple testing. Genes that passed the 5 % FDR threshold were considered to have a true allelic imbalance. In light of our results, we decided to include a control organ in order to compare X/X*-linked allele-specific expression results. We thus performed the same steps as above using kidney transcriptomic data (see supplementary methods). Allele-specific expression analyses were performed on R v4.3.2 (R core Team, 2023).

### g) Gene ontology, candidate genes & enrichment analysis

Enrichment analyses per module were performed using clusterProfiler v4.0 software on R v4.3.2 (R core Team, 2023). We used all genes detected in *M. minutoides* as a background to test for enrichment and applied an adjusted p.value threshold of 0.05 (Benjamini-Hochberg correction for multiple testing) to detect significant enrichments. We also added gene ontology (GO) terms on a gene-by-gene basis for genes whose expression are influenced by the genotype and genes with allelic variations using on the online tool amiGO (https://geneontology.org).

## Acknowledgements

We are grateful to RAM-CECEMA animal facility of University Montpellier and Marie Challe for her help in maintaining the breeding colony. This work was supported by the French National Research Agency (ANR grant SEXREV 18-CE02-0018-01 to F.V.). P.A.S was supported by a postdoctoral fellowship from MUSE program “investissements d’avenir” (ANR-16-IDEX-0006).

