## Supplemental methods, figures and tables for "Influence of gonadal and chromosomal sex on the brain transcriptome in a mouse species with natural sex reversal"

### Supplemental information

### Supplemental methods

##### Sampling, RNA extraction and sequencing of the kidney transcriptome

Kidneys were collected following euthanasia by cervical dislocation of 20 virgin individuals (5 XX, 5 XX*, 5 X*Y and 5 XY; distinct from individuals for brain investigations) aged between 6-9 months old. Samples were kept in RNAlater® (ThermoFisher) for 48h at 4°C and then stored at -80°C until further use. We extracted one kidney randomly chosen (from left or right) following RNeasy® Plus Mini Kit (Qiagen) protocol. For each individual, we grounded the tissue in liquid nitrogen using a mortar and a pestle. We then added lysis buffer (6µl of β-mercapto and 6000µl of RLT Plus buffer per sample) to homogenized shredded materials. RNA was purified following Qiagen provided protocol and total RNA was eluted in 50 µl of RNase free water. Samples were then stored at -80°C until sequencing. Unstranded library preparation and sequencing (Novogen, Cambridge, UK) using NovaSeq 6000 PE150 generated an average of 46 million 150bp paired-end reads. Read quality was assessed using FastQC  v0.11.9. Reads were trimmed for minimal quality (set at 30), minimal length (set at 75bp) and adaptors as well as N bases were removed using Cutadapt v2.8.

##### Read mapping and masking of the X chromosome

In order to decrease mapping bias because we map reads on XX reference genome and because the X and X* have started to stop recombining, we identified SNPs along the X chromosome that are alternatively fixed in our XY and X*Y samples. Those SNPs, called diagnostic SNPs, were then used to mask the reference X chromosome for downstream analyses. Trimmed reads from both kidney and whole brain transcriptomic data were mapped to our XX reference genome (*Mus minutoides* I2396, GenBank accession number GCA_902729485.2) using STAR v2.7.10b with a maximum mismatch tolerance threshold of 6  in 2x150 bp paire-end reads, marking duplicated as well as multi-mapper reads, and sorting reads by coordinates. Bam files from XY and X*Y samples were filtered to keep only read mapping to the X chromosome using samtools (v1.17) view and a bed file containing all scaffolds anchored to *Mus musculus* reference (GRCm39) X chromosome. SNPs were called (separately for brain and kidney samples) in all XY males and X*Y females using GATK (v4.4.0.0) HaplotypeCaller (options -ERC GVCF -ploidy 1). VCF files were joint (per tissue) using GATK GenomicsDBImport and GenotypeGVCFs, and variants were then filtered independently for XY and X*Y individuals using GATK SelectVariants and VariantFiltration. For each called SNP in joint files, samples with depth of coverage lower than three were flagged as not called (--genotype-filter-expression "DP<4" --set-filtered-genotype-to-no-call true), and only positions with total minimum depth of coverage per genotype of 10 (--filter-expression "DP<10"), and at least 3 individuals called per genotype (--filter-expression "AN<3") were considered. For XY samples, variants were filtered to only conserve those in which the reference allele is fixed (--filter-expression "AF>0.1"). For X*Y samples, variants were filtered to only conserve those in which the alternative allele is fixed (--filter-expression "AF<0.9"). Indels, tri-allelic sites and flagged sites were removed with bcftools (v1.17) view (options -M2 -V indels -f PASS). XY and X*Y callsets were merged using bedtools (v2.30.0) intersect, identifying 409 diagnostic SNPs using the brain transcriptome and 316 diagnostic SNPs using the kidney transcriptome. Brain and kidney SNP sets were concatenated using bcftools concat, resulting in a total of 529 diagnostic SNPs that were used to mask the reference XX genome using GATK FastaAlternateReferenceMaker. Trimmed reads were then remapped on XX reference genome masked for X/X* fixed SNPs with the same parameters as above.

**X/X* chromosome inactivation in the kidney**

X/X* allelic counts were retrieved from BAM and VCF files above using the ASEReadCounter tool from GATK v4.4.0. We filtered allelic read counts for mapping quality (≥ 10) , base quality (≥ 10) and the minimum number of bases that passed filters (≥ 10). The ASEReadCounter tool returns allelic counts at the SNP level, we therefore retrieved allelic counts at the gene level using the

ASEReadCounter* pipeline (Mendelevich et al., 2020), which interpolates SNPs and genes' position on genomic scaffolds. Prior to imbalance analyses, we removed genes for which an X* allele was found in XX females or conversely, when an X allele was found in X*Y females (i.e., misdiagnosed SNP: shared polymorphism between chromosomes). We also filtered genes with a minimal count of 10 reads in at least 5 allelic counts out of the 10 (2 alleles x 5 sample replicates). Genes with allelic variation were then averaged between XX* females and plotted to evaluate X/X* chromosome inactivation.

### Supplemental Figures

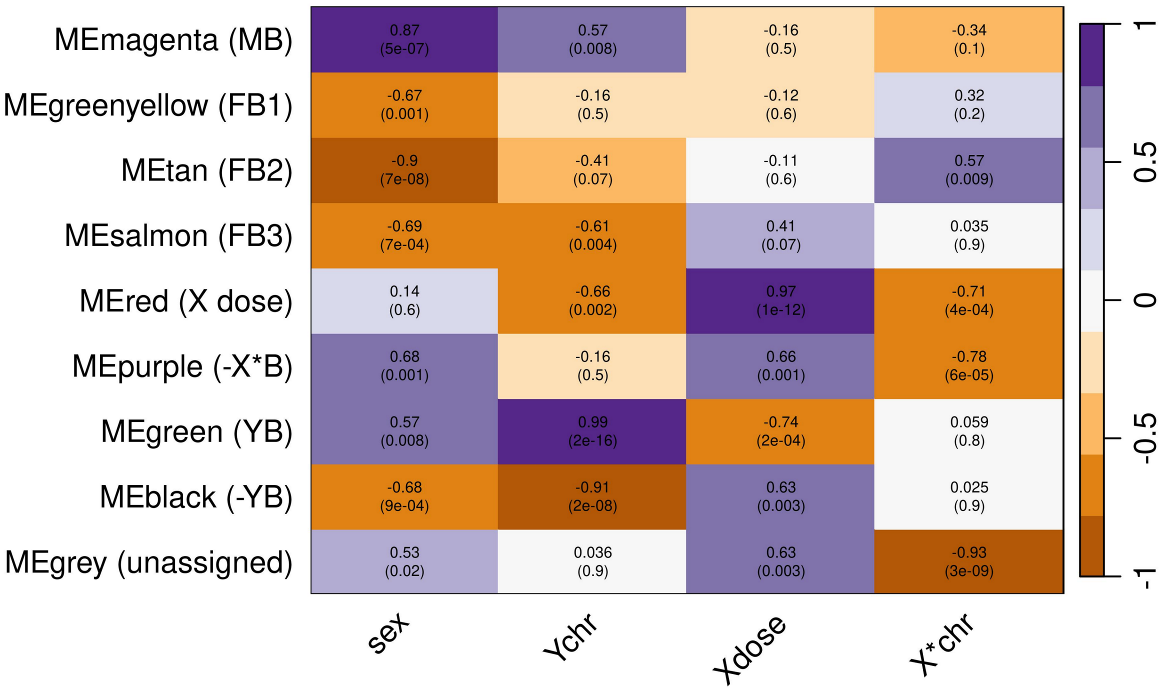


**Figure S1**. Module-biological trait correlations. Each 'MEcolor' correspond to module eigengenes of co-expressed gene modules (i.e., clusters) identified in the clustering analysis. Biological traits represented on the abscissa are : 'sex' : female or male (XX=0 XX*=0, X*Y =0, XY=1); 'Ychr': presence of the Y chromosome / one copy of the X(*) chromosome (XX=0, XX*=0, X*Y =1, XY=1) ; 'Xdose' : X chromosome copy number (XX=2, XX*=1, X*Y =1, XY=0) ; 'X*chr' : presence of the X* chromosome (XX=0, XX*=1, X*Y =1, XY=0). Values represent pearson correlations coefficient ranging from -1 to 1 and calculated from WGCNA software. P.values are shown in brackets. Significance of the correlation allow to detect which traits mainly drive expression patterns in each modules. Correspondence between module colors and cluster names used in this study are shown in supplementary material S1. Turquoise = male-biased genes (MB), pink, blue and greenyellow = female-biased genes (FB1, FB2 and FB3 respectively) ; black = underexpressed in individuals with a single copy of the X chromosome (X dose) ; purple = down-regulation in individuals with an X* chromosome (-X*B) ; red and green = under- and overexpressed genes in individuals carrying a Y chromosome (-YB and YB, respectively).

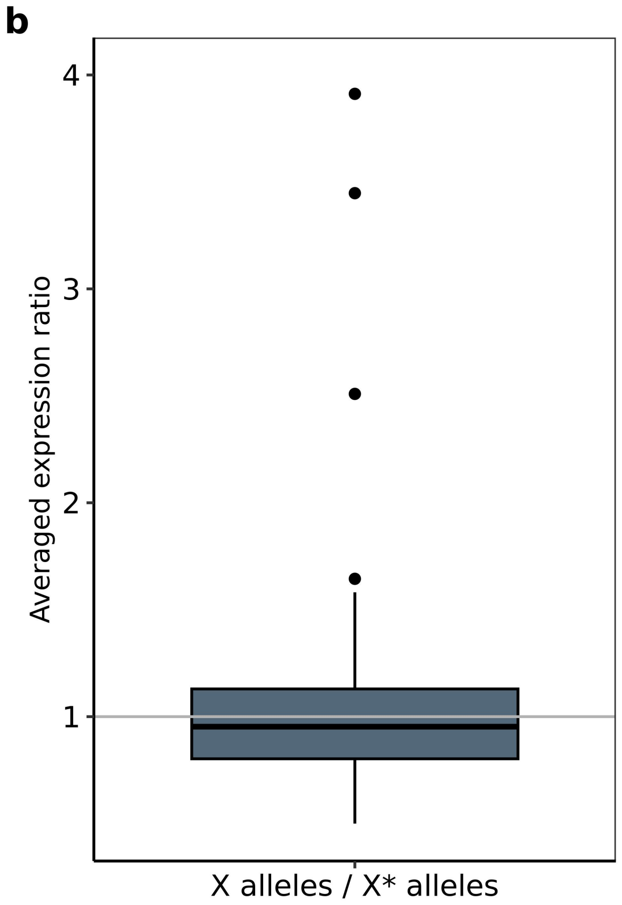


**Figure S2. Averaged allele-specific expression in the kidney of XX*** The boxplot represents ratios of X alleles over X* alleles of genes. Each gene is averaged across individuals. The grey line indicates a ratio of 1 with equal expression of both alleles. All genes showing allelic variation are represented. One outlier individual was removed (see fig. S3, allelic expression ratios per XX* individuals). n=77 genes.

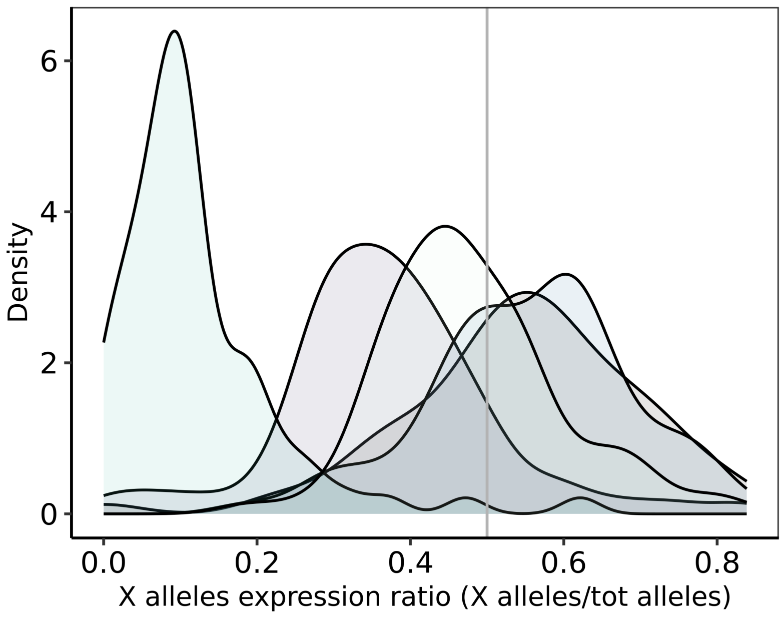


**Figure S3. X/X*-linked allele-specific expression in the kidney transcriptome.** Distribution of X alleles in the five XX* females. Each colour correspond to a specific female. Abscissa values are ratios of X alleles over all alleles per gene. Grey lines indicate a ratio of 0.5 and thus equal expression of both alleles. Values above 0.5 indicate a greater expression of X alleles and conversely, values below 0.5 indicate a greater expression of X* alleles. One outlier individual mostly expressing X* alleles was removed to assess the global pattern of allele-specific expression. $n_{X/X*}$=77.

### Supplemental Tables

| | Autosomes | | X/X* | | | Neo-sex chromosomes | | |
|---|---|---|---|---|---|---|---|---|
| Clusters | N (Freq$_{obs}$) | Permutation test p.value | N (Freq$_{obs}$) | Pearson $\chi^2$ p.value | Permutation test p.value | N (Freq$_{obs}$) | Pearson $\chi^2$ p.value | Permutation test p.value |
| FB | 241 (0.02) | 0.23 | 14 (0.02) | 0.39 | 0.32 | 49 (0.02) | 0.86 | 0.9 |
| MB | 563 (0.05) | 0.90 | 28 (0.05) | 0.54 | 0.69 | 147 (0.05) | 0.22 | 0.27 |
| X dose | 13 (0.001) | 1 | 6 (0.01) | **<0.0001*** ** | **0.01*** ** | 25 (0.009) | **<0.0001*** ** | **0.001*** ** |
| -X*B | 11 (0.0009) | 1 | 4 (0.007) | **0.0005*** ** | **0.01*** ** | 7 (0.003) | **0.04*** | 0.10 |
| YB | 32 (0.003) | 1 | 1 (0.002) | 0.54 | 0.93 | 32 (0.01) | **<0.0001*** ** | **0.001*** ** |
| -YB | 87 (0.008) | 1 | 15 (0.03) | **<0.0001*** ** | **0.001*** ** | 36 (0.01) | **0.004*** ** | **0.04*** |
| Unbiased | 10,068 (0.91) | - | 500 (0.88) | - | - | 2,378 (0.89) | - | - |

**Table S1. Distribution of DE genes and enrichment analysis.** DE gene observed frequencies (Freq$_{obs}$) on sex and neo-sex chromosomes were compared to that of autosomes using Pearson $\chi^2$ tests (one sided hypothesis). Enrichment on sex and neo-sex chromosomes were also tested using permutation tests. We considered regions enriched in DE genes when significance thresholds were passed in both methods. The three female-biased clusters were grouped together. Asterisks indicate the significance level.*** p≤0.001, **p≤0.01,*p≤0.05. Enrichment results were consistent across methods and showed an enrichment in DE genes with a genotype effect on sex and neo-sex chromosomes. MB= male-biased ; FB= female-biased; X dose = genes overexpressed in XX females, followed by XX* and XY individuals, and then X*Y; -X*B= genes underexpressed in X*-carrying females; YB= genes overexpressed in Y-carrying individuals; -YB= genes underexpressed in Y-carrying individuals.

### Supplementary material S1-S2

#### Supplementary material S1

$Log_2FC$ for the 1,362 differential expressed genes (DEG) at least between two genotypes in the African pygmy mouse (edgeR results). Results are based on multiple pairwise comparison (4 genotypes, hence six comparisons), with a FDR threshold at 0.05. Clusters in which genes were placed by the WGCNA software are included as well as corresponding name used in the main text (based on module-trait correlation, see fig. s1). Bold values indicate an absolute FC greater or equal 1.5. Lines highlighted in yellow represent candidate genes at the basis of phenotypes (*) or likely to escape inactivation (†). Enrichment in GO terms in the male-biased cluster and enrichment in GO terms in the FB cluster are shown in different sheets. The top20 GO terms are represented. GO of genes in clusters showing a genotype effect are shown in a different sheet as well.

#### Supplementary material S2

Allelic imbalance results in XX* females. All genes showing allelic variation (i.e., diagnostic SNPs in coding sequence between gametologs) are represented. Values represent results from post-hoc comparisons of allelic expression (following the negative binomial model). Clusters in which genes have been attributed in the WGCNA analysis (see supplemental material S1) are shown. Genes showing a true allelic imbalance are in bold and ontologies of these genes are represented in the following sheet.
